# Assessing protein dynamics on low complexity single-strand DNA curtains

**DOI:** 10.1101/376798

**Authors:** Jeffrey M. Schaub, Hongshan Zhang, Michael M. Soniat, Ilya J. Finkelstein

## Abstract

Single-stranded DNA (ssDNA) is a critical intermediate in all DNA transactions. As ssDNA is more flexible than double-stranded (ds)DNA, interactions with ssDNA-binding proteins (SSBs) may significantly compact or elongate the ssDNA molecule. Here, we develop and characterize low-complexity ssDNA curtains, a high-throughput single-molecule assay to simultaneously monitor protein binding and correlated ssDNA length changes on supported lipid bilayers. Low-complexity ssDNA is generated via rolling circle replication of short synthetic oligonucleotides, permitting control over the sequence composition and secondary structure-forming propensity. One end of the ssDNA is functionalized with a biotin, while the second is fluorescently labeled to track the overall DNA length. Arrays of ssDNA molecules are organized at microfabricated barriers for high-throughput single-molecule imaging. Using this assay, we demonstrate that *E. coli* SSB drastically and reversibly compacts ssDNA templates upon changes in NaCl concentration. We also examine the interactions between a phosphomimetic RPA and ssDNA. Our results indicate that RPA-ssDNA interactions are not significantly altered by these modifications. We anticipate low-complexity ssDNA curtains will be broadly useful for single-molecule studies of ssDNA-binding proteins involved in DNA replication, transcription and repair.

## Introduction

Single-stranded DNA (ssDNA) is a key intermediate in nearly all aspects of DNA metabolism, including transcription, replication, and repair. Cellular ssDNA is rapidly bound by endogenous single-strand DNA binding proteins (SSBs). SSBs rapidly diffuse and stabilize ssDNA from damage and cellular nucleases.^1,2^ SSBs also physically interact with multiple DNA-processing proteins and thus act as recruitment hubs for downstream replication and repair.^3,4^ SSBs bind ssDNA via one or more oligonucleotide/oligosaccharide binding (OB) folds. For example, the homotetrameric *E. coli* single-stranded DNA binding protein (EcSSB) encodes four identical OB-folds that can nonetheless interact with ssDNA in a variety of conformations.^5^ The biological consequences of these alternative binding conformations are still not entirely clear but have been proposed to affect the ssDNA topology and potentially impact EcSSB function.^5,6^

Replication protein A (RPA) is the most abundant eukaryotic ssDNA-binding protein. RPA encodes six OB folds amongst its three heterotrimeric subunits.^7,8^ RPA’s OB folds have a variable affinity for ssDNA, with the strongest interactions encoded in the RPA70 subunit.^9^ RPA binds ssDNA with sub-nanomolar affinities yet is readily displaced by other proteins, such as RAD51.^10–12^ In addition, RPA is hyperphosphorylated in response to DNA damage.^13,14^ RPA phosphorylation is crucial for cell recovery from the DNA damage response in S phase.^15^ Post-translational RPA modifications have been proposed to alter the RPA structure and its interactions with DNA, but the impact of these modifications on ssDNA geometry have been controversial.^16–19^ Here, we use single-molecule fluorescence imaging to probe how EcSSB and human RPA compact ssDNA.

Single-molecule studies of protein-ssDNA interactions have traditionally used two strategies for generating ssDNA substrates. One approach denatures a dsDNA template via mechanical unzipping or addition of a chemical denaturant.^20,21^ Mechanical unfolding of individual molecules is time-consuming and requires a dedicated optical or magnetic tweezer apparatus. Chemical denaturation is frequently incomplete; segments of the template DNA are not melted or can re-hybridize after the denaturant is diluted or washed out of the flow cell. Moreover, harsh chemical denaturants such as NaOH can partially hydrolyze the phosphate backbone. The second strategy generates ssDNA via rolling circle replication (RCR) by a strand-displacing DNA polymerase. This strategy can generate >50,000 nucleotide (nt) ssDNA substrates, especially when strand displacement synthesis is catalyzed by the highly-processive phi29 DNA polymerase (DNAP).^22,23^ RCR products can also be synthesized from short oligonucleotide (oligo) DNA templates.^24,25^ However, fluorescent labeling of the RCR products on the 3’-end is challenging because the ssDNA molecules are highly repetitive. Both approaches also produce complex ssDNA that may encode sequence and secondary structures that confound single-molecule studies. ssDNA templates that contain additional secondary structures must be extended by force, denaturants, or secondary structure-destabilizing proteins.^12,20,21^ These limitations of existing approaches have motivated us to generalize RCR for high-throughput single-molecule DNA curtains.

Here, we describe a method for assembling low-complexity ssDNA curtains via RCR from short oligos. Low complexity ssDNA is largely devoid of standard Watson-Crick base-pairing, resulting in extended ssDNA at minimal applied forces. RCR with phi29 DNAP produces ssDNA molecules exceeding 10^5^ nt in length but is highly dependent on the enzymatic activity of the DNA polymerase. We thus also optimized expression of phi29 DNAP, ensuring highly reproducible RCR products. We describe an improved labeling strategy for specific functionalization of both ssDNA ends. Finally, each ssDNA molecule can be readily converted to double-stranded DNA (dsDNA), allowing accurate measurement of the overall DNA length in nucleotides. DNA curtains organize hundreds of individual ssDNA molecules into arrays on the surface of a microfluidic flow cell for high throughput single-molecule imaging. We validate our strategy by examining the ssDNA length changes induced by different ssDNA-binding modes of the EcSSB. Finally, we utilize this strategy to interrogate whether phosphomimetic human RPA (pmRPA) undergoes significant changes in its ssDNA-binding properties relative to wt RPA. In sum, low-complexity ssDNA curtains will be useful to the broader scientific community interested in the physical changes of ssDNA upon interaction with proteins and other nucleic acids.

## Experimental Section

### Synthesis and assembly of low-complexity ssDNA curtains

PAGE-purified oligos were purchased from IDT. ssDNA circles were prepared by annealing 5 μM phosphorylated template oligo (/5Phos/AG GAG AAA AAG AAA AAA AGA AAA GAA GG) and 4.5 μM biotinylated primer oligo (5/Biosg/TC TCC TCC TTC T) in 1x T4 Ligase Reaction Buffer (NEB B0202S).^24,25^ Oligos were heated to 75°C for five minutes and cooled to 4°C at a rate of −1°C min^−1^. Anneal circles are ligated with the addition of 1 μL of T4 DNA Ligase (NEB M0202S) at room temperature for ~5 hours. Ligated circles can be stored at 4°C for up to a month. Low-complexity ssDNA was synthesized in 1x phi29 DNA Polymerase Reaction Buffer (NEB M0269S), 500 μM dCTP and dTTP (NEB N0446S), 0.2 mg mL^−1^ BSA (NEB B9000S), 10 nM annealed circles and 100 nM phi29 DNAP (purified in-house). The reaction was mixed and immediately injected on the flow cell and incubated at 30°C for 20 minutes. For digoxigenin incorporation as an end-label, 100 μM dTTP was included at a 5, 50, or 500 molar excess over digoxigenin-11-ddUTP (Roche). ssDNA synthesis was quenched by removing excess nucleotides and polymerase with BSA buffer (40 mM Tris-HCl pH 8.0, 1 mM MgCl2, 1 mM DTT and 0.2 mg mL^−1^ BSA).

### Single-Molecule Microscopy

Flow cells were prepared as previously described.^26,27^ Briefly, a 4 mm wide and 100 μm high flow channel is constructed between a glass cover slip (VWR 48393 059) and a custom-made flow cell containing 1-2 μm wide chromium barriers using two-sided tape (3M 665). All experiments were conducted at 37°C under a flow rate of 1 mL min^−1^. Single-molecule fluorescent images were recorded on a prism TIRF microscopy based inverted Nikon Ti-E microscope. The sample was illuminated with a 488-nm laser (Coherent Sapphire) and a 637-nm laser (Coherent OBIS) split by a 638-nm dichroic beam splitter (Chroma). Two-color imaging was recorded using dual electron-multiplying charge-coupled device (EMCCD) cameras (Andor iXon DU897). Subsequent images were exported as uncompressed TIFF stacks and further analyzed in FIJI.^28^

### ssDNA End Labeling

For digoxigenin incorporated ssDNA, ends were labeled with a rabbit anti-digoxigenin primary antibody (Thermo 9H27L19) followed by incubation with an ATTO488-labeled goat anti-rabbit secondary antibody (Sigma 18772). For dsDNA RCR-circle labeled ssDNA, ends were labeled with a mouse anti-dsDNA primary antibody (Thermo MA1-35346) followed by incubation with an ATTO647N-labeled goat anti-mouse secondary antibody (Sigma 50185). End labeling percentage was calculated by counting the number of ssDNA molecules stained with RPA-GFP that were also fluorescently end-labeled divided by total DNA molecules visible in a field of view.

### ssDNA to dsDNA Length Conversion

DNA ends were tracked using custom written FIJI scripts. Briefly, the fluorescent intensity of the ssDNA end label is fit to a two-dimensional (2D) Gaussian function for sub-pixel particle localization. The 2D Gaussian fit is then plotted labeled as a function of the frame number (time). The plateaus in length change are then averaged together to reveal the corresponding length.

Initial ssDNA length was measured at a 1.0 mL min^−1^ flow rate. The ssDNA was converted to dsDNA by injection with 100 nM complementary oligo (at a ratio of 99:1 complementary (5/AGG AGA AAA AGA AAA AAA GAA AAG AAG G, IDT) and ATTO647N-complementary oligo (5/atto647N/AG GAG AAA AAG AAA AAA AGA AAA GAA GG, IBA)). A mixture of unlabeled and ATTO647N-labeled complementary oligo was used to limit laser-induced damage to the ssDNA.

### E. coli SSB Purification

Plasmids overexpressing either EcSSB-GFP (pIF109) or wt EcSSB (pIF122) were transformed into BL21 (DE3) (Novagen) *E. coli* cells. 6L of LB was grown to an OD600 of 0.6 and subsequently induced with 0.5 mM IPTG overnight at 18°C.^12,29^ The resulting pellet was resuspended in Lysis Buffer (25 mM Tris-HCl [pH 7.4], 500 mM NaCl, 1 mM EDTA and 5% (v/v) glycerol, supplemented with 1x HALT protease cocktail (Thermo-Fisher) and 1 mM phenylmethanesulfonyl fluoride (PMSF, Sigma-Aldrich)). Cells were sonicated and centrifuged at 35,000 RCF for 45 minutes. Cellular supernatant was loaded on 5 mL (bed volume) of StrepTactin Superflow resin (IBA Life Sciences) pre-equilibrated with Lysis Buffer. SSB was eluted in 20 mL of Elution Buffer (Lysis Buffer supplemented with 5 mM desthiobiotin (Sigma-Aldrich D1411)) and collected in 5 mL fractions. The elution was pooled and concentrated using spin concentration (Amicon). SSB was dialyzed into Storage Buffer (50 mM Tris-HCl [pH 7.4], 300 mM NaCl and 50% (v/v) glycerol) and flash frozen in liquid nitrogen and stored at −80°C. Protein concentrations were determined by comparison to a BSA standard curve using SDS-PAGE.

### Human RPA purification

The plasmids for wt RPA (pIF47) and RPA-GFP (pIF48) were a generous gift of Dr. Marc Wold and Dr. Mauro Modesti, respectively. The plasmids for pmRPA (pIF430) and pmRPA-GFP (pIF429) was mutagenized using Q5 Site-Directed mutagenesis (NEB E0554S) to create the S8D, S11D, S12D, S13D, T21D, S23D, S29D and S33D mutations of RPA2 from pIF47 and pIF48 overexpression plasmids, respectively. RPA constructs were purified as previously described.^30^

### phi29 DNA polymerase Purification

The gene encoding phi29 DNAP was cloned into a pET19 vector with an N-terminal His-TwinStep-SUMO tag to generate pIF376. The plasmid was transformed into Rosetta (DE3)pLysS (Novagen, 70956) *E. coli* cells. 1L of LB was grown to an OD600 of 0.6 and subsequently induced with 0.5 mM IPTG overnight at 16°C. The resulting pellet was resuspended in Lysis Buffer (25 mM Tris-HCl [pH 7.5], 500 mM NaCl and 5% (v/v) glycerol, supplemented with 1x HALT protease cocktail (Thermo-Fisher) and 1 mM phenylmethanesulfonyl fluoride (PMSF, Sigma-Aldrich)). Cells were sonicated and centrifuged at 35,000 RCF for 45 minutes. Cellular supernatant was loaded on 5 mL (bed volume) of StrepTactin Superflow resin (IBA Life Sciences) pre-equilibrated with Lysis Buffer. The resin was subsequently washed with High Salt Wash Buffer (Lysis Buffer supplemented with 1M NaCl) to remove bound DNA from the polymerase. phi29 DNAP was eluted in 20 mL of Elution Buffer (Lysis Buffer supplemented with 5 mM desthiobiotin (Sigma-Aldrich D1411)) and collected in 5 mL fractions. phi29 DNAP was dialyzed into Storage Buffer (10 mM Tris-HCl [pH 7.5], 100 mM KCl, 1 mM DTT, 0.1 mM EDTA and 10% (v/v) glycerol) with 5 μg purified SUMO Protease (homemade stock generated via overexpression from pIF183) overnight. Digested phi29 DNAP was flash frozen in liquid nitrogen and stored at −80°C. Protein concentrations were determined by comparison to a BSA standard curve using SDS-PAGE.

### SSB Buffer Exchange Experiments

*Ec*SSB Buffer Exchange experiments were conducted in either BSA Buffer or BSA Buffer supplemented with 500 mM NaCl. ssDNA ends were labeled prior to the introduction of SSB. 1 nM SSB(-GFP) (monomers) was included in the buffer flow. Prior to each buffer switch, a 100 μL injection of 100 nM SSB(-GFP) (monomers) was injected to ensure complete coverage of the ssDNA molecules. Data was collected with a 100-millisecond exposure taken every 30 seconds to minimize laser-induced photobleaching.

### RPA and pmRPA-GFP Experiments

Human wt RPA and pmRPA experiments were conducted in BSA Buffer supplemented with 150 mM NaCl. ssDNA ends were labeled prior to the introduction of RPA. Initial ssDNA length was measured at a 1.0 mL min^−1^ flow rate. The ssDNA was coated by injection with 5 nM (pm)RPA. RPA and pmRPA-GFP exchange experiments were conducted in BSA Buffer. 10 nM pmRPA-GFP was included in buffer flow to ensure complete coating of the ssDNA molecule. The buffer was then switched to contain 10 nM wt RPA. Data was collected with a 100-millisecond exposure taken every 5 seconds.

## Results and Discussion

For single-molecule imaging, we adapted an oligo-based RCR method for generating low-complexity ssDNA templates (**Fig. 1**).^24,25^ A short 28-nt phosphorylated template and a 12-nt biotinylated primer oligo are annealed to form a mini-circle. The addition of T4 DNA ligase converts the nicked mini-circle into a covalently closed form (**Fig. 1A**). The template strand is designed to eliminate Watson-Crick base-pairing in the synthesized ssDNA molecule.^24,25^ The 5’-end of the minicircle primer is biotinylated for downstream assembly on the surface of a microfluidic flow cell (**Fig. 1B**). The minicircles are introduced into the flow cell and RCR is performed *in situ* with phi29 DNAP. Performing RCR in the flow cell reduces pipettor-induced shearing of the >50,000 nt-long ssDNA molecules and generating longer ssDNA substrates.^31^ The resulting ssDNA molecules consist of tens of thousands of repeating 28 nt sequences. These ssDNAs are organized at microfabricated chromium barriers via the addition of hydrodynamic force (buffer flow; **Fig. 1C**). The free 3’-end of the ssDNA molecule can be fluorescently labeled, allowing simultaneous observation of fluorescent ssDNA-binding proteins and how these proteins alter the ssDNA length. Repeated cycles of toggling buffer flow on and off retract the ssDNA to the chromium barrier, indicating that the ssDNAs are not associated with the flow cell surface (**Fig. 1D**). Hundreds of individual molecules are readily observed in a single field-of-view for rapid data acquisition.

**Figure 1:**
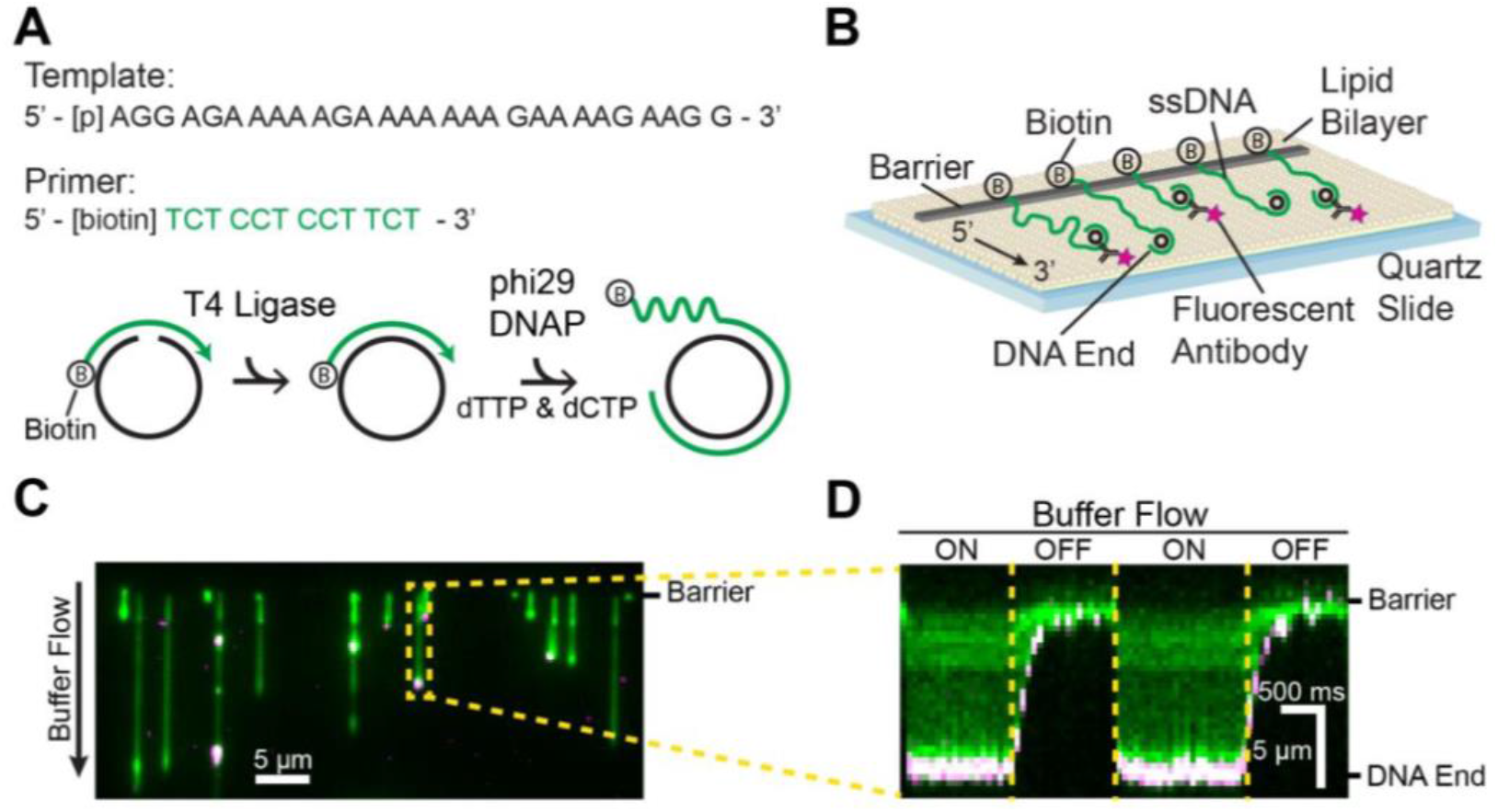
Assembly of low complexity ssDNA curtains. (A) A phosphorylated template (black) and biotinylated primer (green) are annealed and treated with T4 DNA ligase to make minicircles. Low complexity ssDNA comprised solely of thymidine and cytidine is synthesized via rolling circle replication by phi29 DNAP. (B) Low complexity ssDNA curtains with fluorescent end-labeling. The 3’ end of the ssDNA was labeled with a fluorescent antibody. (C) RPA-GFP (green) coated ssDNA with fluorescent end labeling (magenta). (D) Kymograph of a representative ssDNA in panel (C) with buffer flow on and off, indicating that the ssDNA is anchored to the surface via the 5’-biotin tether.

The length of the ssDNA substrates is critically dependent on the enzymatic activity of phi29 DNAP. We found that commercially available enzyme had variable activity and produced short ssDNA substrates (**Fig. S1**). We therefore developed an alternative protocol for rapid purification of phi29 DNAP. Codon-optimized phi29 DNAP was expressed as an N-terminal SUMO-protein fusion and purified via a single gravity-flow StrepTactin column. Following elution from the column with desthiobiotin, the enzyme was dialyzed in the presence of SUMO protease to remove the SUMO domain. Home-made phi29 DNAP was stored at −80°C and retained high RCR activity for >8 months. The homemade construct consistently synthesized ssDNA molecules that were >5-fold longer than commercial polymerases (**Figs. S1C-D**).

RCR generates a broad distribution of ssDNA molecule lengths. We therefore sought to relate the length of each ssDNA molecule in microns to its corresponding length in nucleotides. Our overall strategy is to measure the length of each ssDNA molecule at a given flow rate, followed by conversion of that ssDNA to dsDNA. The well-characterized Worm-Like Chain (WLC) model can then be used to correlate the length of the dsDNA to its corresponding composition in base-pairs. The number of base-pairs in the dsDNA molecule also defines the nucleotide-length of the parental ssDNA molecule.

We first attempted to fluorescently label the 3’-end of the ssDNA molecules via enzymatic incorporation of digoxigenin-11-ddUTP (ddUTP) to terminate ssDNA synthesis. The digoxigenin moiety was fluorescently labeled by anti-digoxigenin conjugated quantum dots, as has been described for dsDNA curtains previously.^32^ However, using high ddUTP to dTTP ratios resulted in very short ssDNA molecules (**Fig. 2A**). The median lengths of ssDNA were 2.3 μm [IQR 1.8-2.6 μm, N=31], 3.7 μm [IQR 2.3-6.0 μm, N=25] and 3.0 μm [IQR 2.0-5.5 μm, N=27] for 1:5, 1:50 and 1:500 ddUTP to dTTP, respectively. Decreasing the ddUTP concentration also decreased the single-molecule labeling efficiency, resulting in ~90% of unlabeled ssDNAs (**Fig. 2B**). We thus pursued an alternative strategy that labels the minicircle dsDNA that remains on the 3’-end of the ssDNA. Using an anti-dsDNA antibody conjugated to a fluorescent dye, we labeled the 3’ ends of ~66% of the ssDNA molecules (N=107/162), while maintaining an overall length of 4.9 μm [IQR 2.9-7.1 μm, N =24]. The anti-dsDNA antibody is highly specific, as there was minimal observable fluorescence on the ssDNA segment of the molecules. This labeling approach should be widely useful to researchers generating ssDNA via RCR.

**Figure 2:**
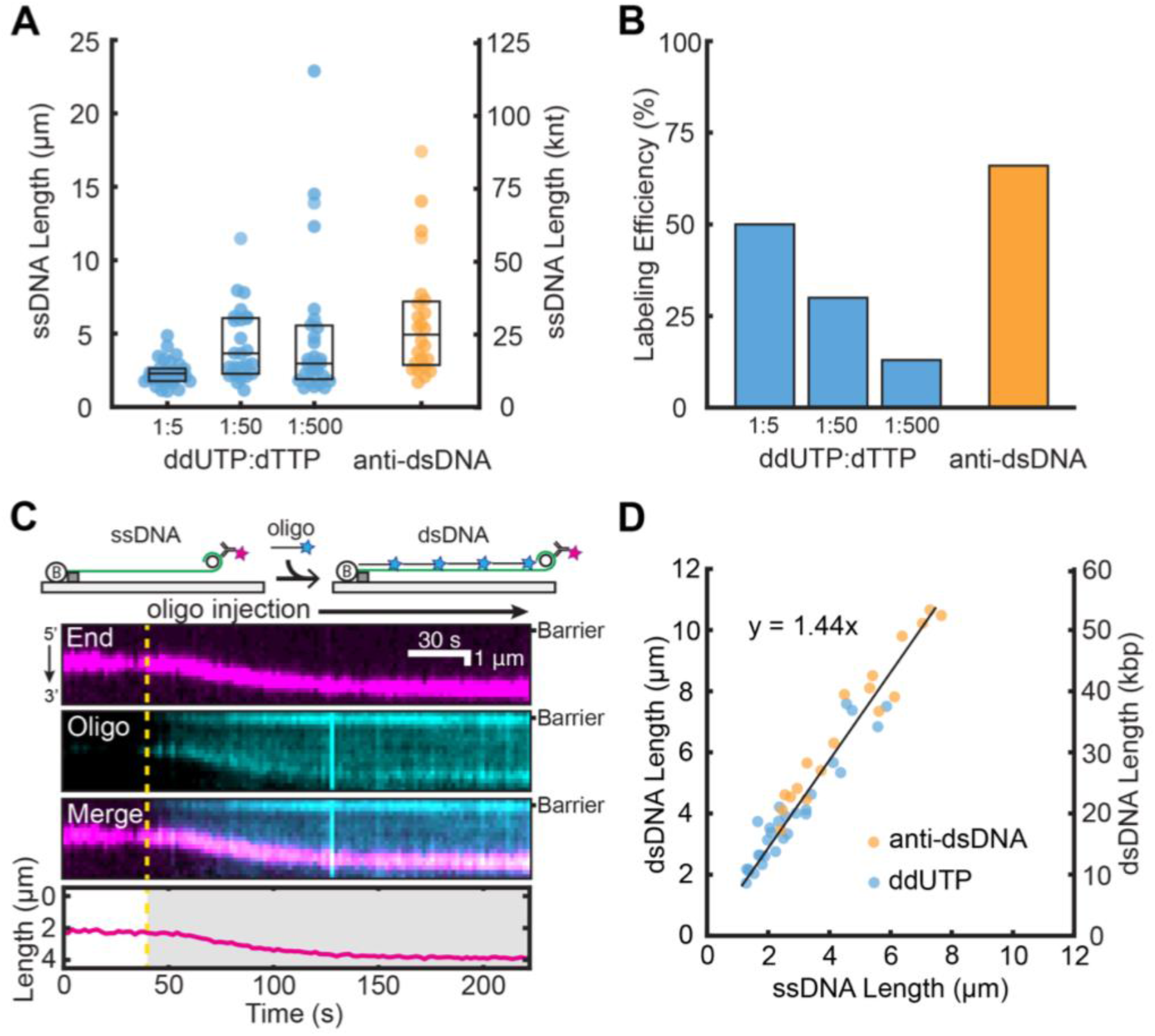
Efficient end labeling and length calibration of low complexity ssDNAs. Comparison of (A) ssDNA lengths and (B) ssDNA end-labeling efficiencies following treatment with ddUTP-dig or fluorescent antibodies. Boxplots denote the first and third quartiles of the data. (C) Cartoon illustration (top) and a representative kymograph of ssDNA (magenta labels the 3’-end) elongating following injection of a complementary oligo (cyan). The DNA length was tracked by fitting the 3’ -end signal (bottom). Dashed line: oligo injection time. (D) The ssDNA to dsDNA length change was quantified by either anti-digoxigenin ddUTP (blue) or anti-dsDNA (orange). Experimental data was fit to a line (black) with slope 1.44 ± 0.05.

Next, we exploited the highly repetitive RCR product to convert the ssDNA molecules into dsDNA by incubation with a complementary oligo (**Fig. 2C**). Injection of a complementary oligo (100 nM final concentration, 1% labeled with ATTO647N) showed an increase in the DNA extension (**Figs. 2C & 2D**). Conversion of ssDNA to dsDNA was also visible as the coating of the ssDNA substrate with ATTO647N signal due to oligo hybridization. The DNA extends and eventually plateaus, indicating a fully-hybridized dsDNA molecule. Comparing the ssDNA length to the final dsDNA extension showed a linear correlation (**Fig. 2D**). The ssDNA to dsDNA extension has a slope of 1.44 ± 0.05 at a flow rate of 1 mL min^−1^. We measured that dsDNA is extended to 0.84 times the length of crystallographic B-form dsDNA (0.29 nm inter-base distance) at this flow rate, corresponding to ~1 pN of applied force (**Fig. S2**). Together, these measurements directly report the size of each ssDNA molecule in nucleotides.

### NaCl induced structural transitions of E. coli single-stranded DNA-binding protein

*Ec*SSB is critical for DNA replication, repair and recombination.^3,33^ It engages ssDNA in various conformations with two predominant footprints of 35- and 65-nucleotides, engaging two or four OB-folds on average, respectively (**Fig. 3A**).^5,34^ These forms exist in equilibrium but interestingly, the ssDNA binding mode of *Ec*SSB is directly impacted by the concentration of cation in solution, with higher concentrations favoring the compacted 65-nt binding mode.^35,36^ The *in vivo* significance of the binding modes is unclear, although the 65-nt binding mode is dispensable for growth.^6^ Previous AFM and single-molecule experiments have interrogated the compaction and extension of EcSSB with differing results.^37,38^ Here, we use ssDNA templates to explore the salt-induced conformational changes of EcSSB on ssDNA.

**Figure 3:**
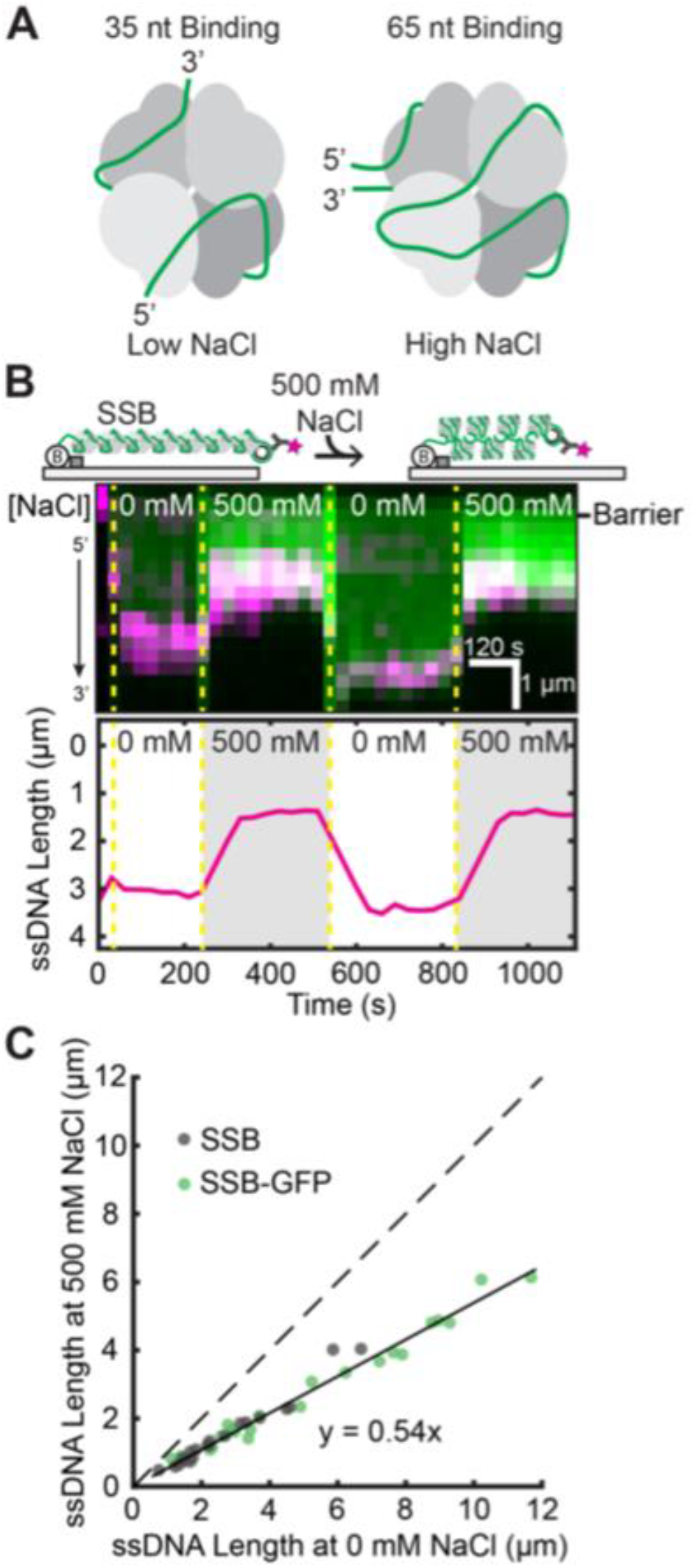
Analysis of EcSSB Binding Modes on ssDNA. (A) Illustration of proposed *Ec*SSB-ssDNA binding modes^5,34^. (B) Cartoon (top) and kymograph of *Ec*SSB-GFP (green) on ssDNA (3’-end in magenta). Dashed yellow lines denote when the buffer switched between high and low NaCl concentration. The ssDNA end position was determined via single-particle tracking (bottom). (C) Correlation between ssDNA-*Ec*SSB length at 0 mM and 500 mM NaCl. Solid black line is a linear fit to the data (N=23 EcSSB-GFP and 20 wt EcSSB). Dashed line represents a slope of 1.

As a proof of principle, we sought to test whether our ssDNA method is able to precisely measure the length of EcSSB conformations. *Ec*SSB-GFP or wt *Ec*SSB were incubated with end-labeled ssDNA substrates at 0 mM and 500 mM NaCl concentrations (**Fig. 2B**). We observed a drastic shortening of the ssDNA when the buffer is switched from 0 mM to 500 mM NaCl (**Fig. 3B**). The shortening is fully reversible for both wt *Ec*SSB and *Ec*SSB-GFP, as the ssDNA re-extends to the initial length when the buffer is returned to 0 mM NaCl. Individual observations of ssDNA length transitions showed a linear fit with a slope of 0.54 ± 0.01, as would be predicted from EcSSB binding in the 35-nt and 65-nt modes (**Fig. 3C**). Moreover, our results indicate that GFP does not impact EcSSB’s NaCl-dependent interactions with ssDNA. These results closely mirror the condensation of ssDNA-EcSSB by AFM, where the 65-nt binding mode length was ~60% that of the extended 35-nt mode.^37^ However, we did not observe higher levels of compaction seen in a previous single-molecule study.^38^ This discrepancy may arise from the technical challenges associated with fully denaturing the 48.5 kbp DNA substrates used in the earlier study. In addition, the prior measurements were conducted at a different force regime (0.1-0.5 pN). Here, we conclude that *EcSSB* compacts the ssDNA in accordance with the number of OB-folds engaged by each DNA-binding mode.

### Conformational binding mechanism of RPA and phosphomimetic RPA

RPA is the most abundant SSB in eukaryotic cells. RPA also serves as a signal for the DNA damage response and stalled replication via hyperphosphorylation on the N-terminus of RPA32.^4,39^ Phosphorylated RPA (pRPA) is excluded from replication centers *in vivo*, highlighting a direct role in regulating replication.^40^ pRPA can regulate the DNA damage response via two possible mechanisms: (i) pRPA acts as a key mediator to recruit DNA damage proteins and/or (ii) pRPA induces structural changes that alter the ssDNA binding modes or properties of RPA.^7,41^ Previous studies have detailed two principle modes of DNA binding by RPA. The first includes DNA binding domains (DBDs) A and B, which occupy a footprint of approximately eight nucleotides.^42^ Secondly, an extended binding footprint of ~30 nucleotides that engages the DBDs A, B, C and D.^43^ Structural examinations of the effects of pRPA on conformational dynamics suggest that the N-terminus and DBD B of RPA70 may interact strongly with the highly phosphorylated N-terminus of RPA32, potentially impacting the binding interaction with ssDNA.^16,41,44^ Indeed, previous studies showed that that pRPA binds with poorer affinity to short eight-nt oligos, hypothesizing a transition to the extended conformation.^41^ However, RPA and pRPA have similar binding affinities towards longer oligos.^16,17^ Therefore, the highly negatively charged N-terminus of RPA32 may be sufficient to alter the DNA-binding properties of RPA.

Previous single-molecule studies have utilized RPA to extend ssDNA molecules.^45,46^ We sought to test whether there is appreciable ssDNA length change between pRPA and RPA using our end-labeled ssDNA assay. In order to assay *in vitro*, we generated a phosphomimetic variant of RPA (pmRPA) by introducing eight mutations in the N-terminus of RPA32 (**Fig. 4A**). This phosphomimetic RPA variant recapitulates pRPA phenotypes in vivo and is indistinguishable from pRPA in vitro.^16,17,40^ We initially observed the change in ssDNA length upon introduction of either RPA or pmRPA (**Fig. 4B**). Interestingly, we do not observe significant differences in the extension of ssDNA from RPA or pmRPA with a combined slope of 1.84 ± 0.06. To reaffirm our observation, we sought to do a protein replacement assay. Initially, ssDNA was bound with pmRPA-GFP and then replaced with RPA. We did not observe any length changes between binding of pmRPA-GFP and RPA on end-labeled ssDNA curtains (**Fig. 4C**). Examination of the pmRPA-GFP fluorescent intensity shows a steep decline upon introduction of RPA, indicating a complete exchange of pmRPA-GFP to RPA, as seen for previous single-molecule RPA experiments.^12^ Analysis of individual molecules shows that the there is a linear correlation between pmRPA-GFP length and RPA length with a slope of 1.0 ± 0.03, confirming the same ssDNA extension for pmRPA and wt RPA (**Fig. 4D**). Several lines of evidence suggested that pmRPA may bind ssDNA in an altered conformation relative to RPA.^16,41,44^ However, our results indicate that pmRPA does not undergo a structural transformation that affects the end-to-end length of ssDNA, at least on extended DNA substrates. We cannot rule out interactions between the N-termini of RPA32 and the RPA70, as reported by several recent studies.^16,41,44^ Our results are minimally consistent with a model where the N-terminus of RPA70 does not interact appreciably with ssDNA and thus does not change the binding mode of the wt RPA complex relative to pmRPA.^39,47^ Taken together, these observations suggest that the major function(s) of pRPA in the cell is likely as a loading and recruitment platform for additional repair proteins to the sites of DNA damage.

**Figure 4:**
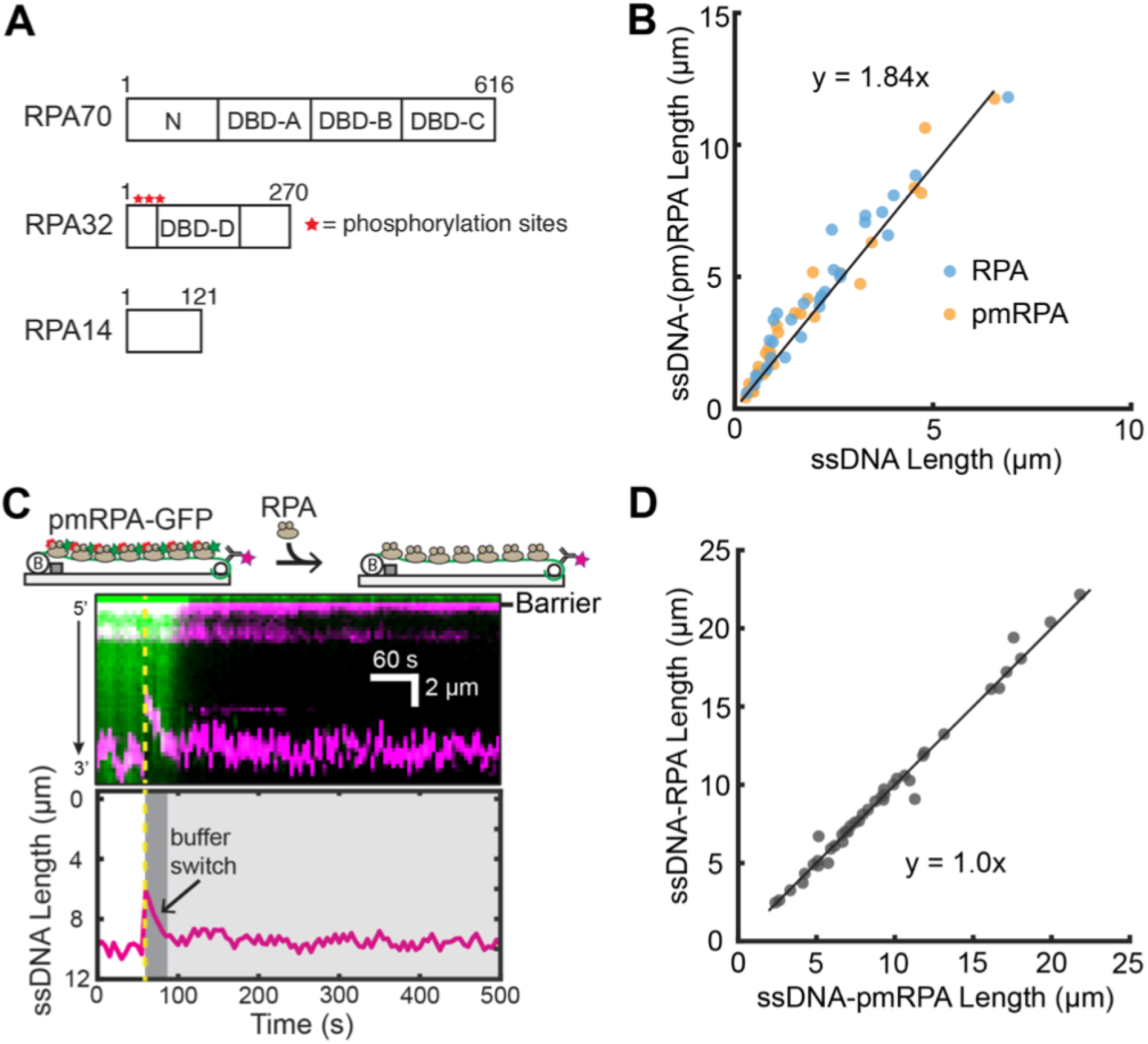
Analysis of RPA Binding Modes on ssDNA. (A) Schematic of the DNA binding domains (DBDs) of the heterotrimeric RPA. Red stars denote the approximate locations of eight phosphorylation sites of RPA2. (B) Both RPA and pmRPA extend the ssDNA to a similar extent. Solid black line is a linear fit to both RPA (N=31 molecules) and pmRPA (N=34 molecules). (C) Cartoon (top) and kymograph of ssDNA-bound pmRPA-GFP (green) as it is replaced by wt RPA (unlabeled). The corresponding change in DNA length is shown in magenta. Dashed yellow line denotes when pmRPA-GFP was replaced with RPA in the buffer. Bottom: ssDNA end-label tracking. (D) Scatter plot of ssDNA length with pmRPA and RPA. Solid black line is a linear analysis of >3 flow cells (N=41 DNA molecules).

## Conclusion

Here, we describe an approach to generate >50,000 nt-long, low-complexity ssDNA curtains via *in situ* RCR. With these templates we are able to estimate the size of the ssDNA in nucleotides by application of buffer flow alone, making these templates highly useful for measuring the effects of ssDNA interacting molecules. In addition, the ssDNA is fluorescently end-labeled allowing dynamic visualization of wild-type, unlabeled single-stranded binding proteins. To increase the adoption of this experimental platform, we describe a few key practical considerations below.

The template and primer oligos can readily be purchased, aliquoted and stored to provide reagents for mini-circle generation for several years. Annealed and ligated mini-circles can be stored at 4°C for up to a month without compromising efficiency. The in-house produced phi29 DNAP yields significantly longer ssDNA molecules and is active for months when stored at −80°C. In addition, performing RCR *in situ* results in greater ssDNA end-labeling, likely by reducing DNA shearing during DNA curtain assembly.^31^ In summary, we describe a method for assembling low-complexity ssDNA substrates for high-throughput single-molecule studies. Because this ssDNA lacks Watson-Crick base-pairing propensity, it is more equivalent to poly-T oligos that are frequently used in ensemble biochemical experiments. This method will be widely applicable to other single-molecule observations of protein-nucleic acid interactions.

## Acknowledgments

We are indebted to Dr. Marc Wold and Dr. Mauro Modesti for sharing over-expression vectors, and to members of the Finkelstein lab for careful reading this manuscript. This work is supported by the NSF (Grant 1453358 awarded to I.J.F). and by the Welch Foundation (Grant F-1808 awarded to I.J.F.). Michael Soniat is supported by a Postdoctoral Fellowship, PF-17-169-01-DMC, from the American Cancer Society. The content is solely the responsibility of the authors and does not necessarily represent the official views of the National Institute of General Medical Sciences or the National Institutes of Health.

The authors declare no conflict of interest.

This article contains supporting information online.

## Author Contributions

J.M.S., H.Z. and I.J.F designed the experiments and wrote the manuscript with input from all co-authors. J.M.S. and H.Z. performed the experiments. M.M.S. prepared the RPA and pmRPA constructs. All authors have reviewed and approved to the submitted manuscript.

